# Phosphorylation of UBE2J1 at serine residue S184 contributes towards infection and cellular syncytialization by Vesicular Stomatitis Virus

**DOI:** 10.64898/2026.04.12.717905

**Authors:** Noor D. Algoufi, Emily B. Walsh, Zakiyah I. Fallata, Sawsan S. Alamri, Anwar M. Hashem, John V. Fleming

## Abstract

The ubiquitin-conjugating enzyme UBE2J1 functions in the proteasomal degradation of proteins at the ER. Existing evidence suggests that it plays a role during viral infection, with elevated UBE2J1 levels generally associated with increased infection. This is particularly relevant for some RNA viruses; however, the regulation of UBE2J1 during infection has not been well studied. Here, we used a BHK21 cell model to demonstrate that UBE2J1 overexpression promotes the replication of Vesicular Stomatitis Virus (VSV), as indicated by a significant increase in viral titres. To better understand the underlying molecular processes, cells were co-transfected to express the VSV-G protein and wild-type UBE2J1 protein, and we observed a significant increase in the syncytial fusion area. This effect was not observed when catalytically inactive (C91S) or phospho-deficient (S184A) forms of the protein were used. Interestingly, overexpression of a truncated, non-ER localized form of UBE2J1 (ΔTM) led to the largest increase in the syncytial fusion area. This arose as a result of increased syncytia size, and may indicate an enhanced cellular role if soluble forms of UBE2J1 are not anchored to the ER. Additional studies using truncated, mutated and wild-type proteins confirmed that UBE2J1 increases VSV viral replication and is associated with an increase in the number of infection plaques. Considering the emerging evidence for UBE2J1 involvement in viral infection, our finding should help in understanding the role of this protein in viral pathogenesis and cellular processes linked to syncytialization.

## INTRODUCTION

UBE2J1 (Ubc6e) is a ubiquitin-conjugating enzyme that functions in the ubiquitination and proteasomal degradation of misfolded proteins. The enzyme is localised in the endoplasmic reticulum (ER), where membrane anchorage is mediated by carboxyl-terminal sequences (amino acids 288-318) (1). The amino-terminal region faces the cytoplasm, where it interacts with translocon components that promote the retro-translocation of misfolded proteins through the ER membrane and catalyses their ubiquitination and subsequent degradation by cytosolic proteasomes(2, 3).

Although UBE2J1 was originally described to be involved in the Unfolded Protein Response (UPR) and the process of ER-associated degradation (ERAD) (1, 4), it has since been implicated in several cellular processes, including immune function. For example, it contributes to TNF-α signalling (5) and the regulation of NF-kB (6). In the context of immune signalling, p38 MAPK and lipopolysaccharide have been shown to promote the phosphorylation of UBE2J1 at serine 184 (5). Although the cellular consequences of this phosphorylation are still being characterised, it appears to be instrumental in the re-establishment of cellular homeostasis following ER-associated stresses (7).

A central role for UBE2J1 during viral infection is also emerging. At one level, it is involved along with HRD1 (HMG-CoA Reductase Degradation protein 1/ Synoviolin) in the processing of antigens for MHC-class I presentation (2, 8). At another level, several studies have demonstrated a direct contributory role in viral propagation, with a positive correlation generally observed between viral infection and UBE2J1 levels (9-12). For several RNA viruses this is linked to the ability of UBE2J1 to disrupt antiviral response networks, including promoting the degradation of Interferon Regulatory Factor (IRF) family members, and subsequent reduction in the production of antiviral interferons (13).

Initial observations describing this mechanism were made with Dengue virus (DENV) where UBE2J1 promotes the ubiquitination and proteasomal degradation of IRF3 (13); however, the paradigm has since been extended to include subtype A avian leukosis virus where it disrupts the STAT3/IRF1 pathway (14), and for Spring viremia of carp virus, where it promotes the ubiquitination and degradation of IRF7 (15). The chicken UBE2J1 protein has been shown to be important for avian leukosis virus (subtype A) infection, with a S184A mutant being less capable of regulating levels of STAT3 (14). However, it remains unclear whether UBE2J1 phosphorylation is actively regulated during infection.

For the SARS-CoV-2 virus, which is also an RNA virus, transcriptomic analysis suggests disruption of ubiquitin-proteasome pathway components in infected patients, including changes observed specifically in UBE2J1 transcript levels (9). It has also been noted that SARS-CoV-2 infection is associated with extensive syncytialization, which involves the fusion of cell membranes to form large multinucleated cells (16). Although the formation of these giant cells is a feature of normal developmental processes, it is often subverted and exploited by infecting viruses (17), where the large fusion areas can contribute towards certain stages of infection (16). Although syncytialization can be linked at the cellular level to the interferon regulatory pathways (18) it has not previously been linked to UBE2J1 expression.

The Vesicular Stomatitis Virus (VSV) is a single-stranded negative-sense virus, which is frequently used as an experimental model for RNA virus infections. Infection is mediated by the VSV glycoprotein (VSV-G), which is located on the surface of the virion and is believed to bind to the low-density lipoprotein receptor (LDL-receptor) on target human cells (19). Subsequently, once the viral genome is expressed and VSV-G is located on the surface of infected cells, the interaction of virion-expressed G-protein with infected cell-expressed G-protein promotes and strengthens the infection and syncytia formation (20). On its own, and in the absence of the virus, transfection of certain cell types with VSV-G can promote cellular processes associated with viral infection, including the formation of syncytia (21).

Here, we assessed the involvement of UBE2J1 in VSV infection. Using BHK21 cells that were either infected with the virus or transiently transfected with the VSV-G protein, we demonstrate a role for UBE2J1 in syncytialization and highlight the importance of serine S184 in promoting infection.

## MATERIALS AND METHODS

### Cell culture

HEK293T (ATCC) and BHK21 (ATCC) were cultured in Dulbecco’s Modified Eagles Medium (DMEM) supplemented with 10% or 5% Fetal Bovine Serum (FBS) respectively and 1% Penicillin/Streptomycin (100 IU/ml) (Sigma) in a humidified incubator at 37°C with 5% CO_2_.

### Plasmids

The coding sequence for human UBE2J1 (amino acids 1 through 318) was PCR amplified from a human testes cDNA library using sense (5’ CTC AAG CTT TCG AAT TCT GCA GGG ATG GAG ACC CGC TAC AAC 3’) and antisense (5’ TGG GTA CGT CGA CAA CTC AAA GTC AAA TAT GTA TTC GTT TGC CAG 3’) primers and cloned sequentially into the Pst1 and Sal1 sites of pEP7-FL vector (22, 23) using In-Fusion HD cloning (Takara) and then subsequently into the PstI and SalI sites of pEP7-His vector following restriction digestion and T4 ligation (Thermo Fisher) to give pEP7-UBE2J1-WT-His. Following PCR amplification using sense (5’ CTC AAG CTT TCG AAT TCT GCA GGG ATG GAG ACC CGC TAC AAC 3’) and antisense (5’ ATG ATG ATG ATG ATG CGT CGA CCC ACC ATG ATC AGT GTG GTT GTC TCT 3’) primers, carboxy-truncated (amino acids 1 through 287) coding sequences were cloned into the pEP7-His tagged vector giving rise to the pEP7-UBE2J1-ΔTM-His vector. The His-tagged plasmid encoding the full-length cDNA (pEP7-UBE2J1-WT-His) was used as template for Quikchange PCR mutagenesis to generate the pEP7-UBE2J1-C91S-His and pEP7-UBE2J1-S184A-His plasmids using sense (C91S; 5’ GTG GGC AAG AAA ATC GCT TTG AGC ATC TCA GGC 3’ and S184A; 5’ CTG GCT AGG CAA ATC GCC TTT AAG GCA GAA GCT 3’) and complementary antisense primers. When appropriate UBE2J1 expressing plasmids were transfected with empty vector control (pcDNA3.1) or pCAGGS-G-Kan (Kerafast) expressing the VSV-G protein. For HEK293T cells, plates were coated with Poly-L-Lysine. For transfections, cells were transfected with Lipofectamine 2000 (Invitrogen) as advised by the manufacturer. OptiMEM (Thermo Fisher) or an equivalent was used as the diluent.

### Western blotting

Cells were harvested in RIPA buffer as described elsewhere (24) and lysates were denatured in Laemmli buffer by 11% SDS-PAGE. Following transfer, membranes were probed with anti-His (Genescript), anti-tubulin (Sigma), anti-VSV-G tag (ThermoFisher), or bespoke anti-UBE2J1 and phospho specific anti-UBE2J1 pSer184 antibodies (Davids Biotechnology).

### Construction and generation of VSV-GFP Virus

The construction and generation of the recombinant VSV-GFP virus was performed as previously described (25). Briefly, the recombinant VSV (rVSV) plasmid was generated by cloning the VSV glycoprotein G (VSV-G) and the GFP gene in the VSVΔG-P/M-MCS2-2.6 expression plasmid (Kerafast). The VSV-G gene was subcloned between MluI and NheI restriction sites between matrix (M) and the RNA-dependent RNA polymerase (L) genes, whereas the GFP was subcloned between AscI and AvrII restriction sites between the phosphoprotein (P) and M proteins. Construction of generated plasmids was confirmed by sequencing and restriction digestion. To rescue the recombinant virus, 70% confluent BHK-21 cells in six 6-well-plates were infected with a recombinant vaccinia virus expressing T7 RNA polymerase at an MOI of 10. After 1.5 h incubation, virus inoculum was removed and cells were transfected with 2 μg of rVSV plasmid, in addition to VSV helper plasmids including N (1 μg), P (1.25 μg), and L (0.25 μg) using Lipofectamine 2000 Transfection Reagent (Thermo Fisher) in Opti-MEM (Thermo Fisher) according to the manufacturer’s instructions. After 2 h incubation at 37 °C, 1 mL of complete growth medium DMEM was added. After 72 h post-transfection at 37 °C, the supernatants containing the rescued virus was collected, centrifuged at 600xg for 5 minutes, filtered twice through 0.22 μm filters to remove the remaining vaccinia. Filtered supernatant was passaged once on BHK-21 cells until cytopathic effects (CPE) and GFP expression were very apparent. Rescued virus was propagated and titrated on BHK-21 cells (See Supplementary materials).

### Virus Infection Conditions

BHK-21 cells were grown in a 12 well plate until ∼ 60-80% confluent. The media was removed, and the cells were washed with warm PBS before adding the virus inoculum diluted to the required multiplicity of infection (MOI) in serum-free DMEM (without supplements). The plate was then incubated in a humidified incubator at 37 °C and 5% CO_2_ and gently rocked every 10 min. After 1 h, the inoculum was removed, the cells were gently washed three times with warm PBS, and then cultured in complete DMEM supplemented with 5% FBS. Culture supernatants were harvested 24 h post-infection (hpi). The supernatants were stored at -80 °C to be used in plaque assay experiments.

### Plaque assay

The titres of purified VSV-GFP virus stocks or our samples were determined by plaque assay on confluent BHK-21 cells. BHK-21 cells were seeded in 6-well plates (1 x 10^6^ cells/well) or 2 x 10^5^ BHK-21 cells (per well of a 12-well tissue culture plate). After 24 h of incubation, a series of 10-fold dilutions of the virus stock was prepared using serum- and supplement-free DMEM, and the virus inoculum replaced the seeding media. The virus was allowed to adsorb at 37°C and 5% CO_2_ for 1 h with rocking the plate every 10 min. After 1 h of incubation, the virus inoculum was removed and overlaid with 2ml medium prepared at a 1:1 ratio with pre-warmed 1.6% Bacto agar (Thermo Fisher) to 2x DMEM (Thermo Fisher) supplemented with penicillin/ streptomycin (final concentration 100 IU/ml - Gibco) and FBS (final concentration 5% -Sigma-Aldrich). Plates were incubated at 37°C and 5% CO_2_ for 36 h, and plaques were counted using a fluorescence microscope or following careful removal of the agarose with a spatula and fixing of the cells by using methanol: glacial acetic acid (1:3 ratio) and staining with crystal violet solution for 30 min. The wells were rinsed three times with water and allowed to dry. Plaques were counted and the titre was calculated as plaque-forming units per ml (PFU/ml).

### Syncytia Formation Assay

In order to analyse VSV-G protein-driven cell-to-cell fusion, BHK-21 cells were grown in 12-well plates and transfected with an expression vector for the VSV-G protein (pCAGGS-G-Kan). In addition, cells transfected with empty plasmid served as control or UBE2J1 plasmids. At 48 h post transfection, the cells were washed with PBS and fixed by incubation (15-20 min, room temperature) with 4% paraformaldehyde solution (Sigma). Thereafter, the cells were washed with PBS, then washed with water to remove salts, air-dried and stained with hematoxylin gill II solution (10 min, room temperature; Surgipath® (Leica Biosystems)). Next, cells were washed three times with 1x Scott’s solution (from 10 x solution containing MgSO_4_7H2O 200 g/l and NaHCO_3_ 20 g/l), plates were placed in double distilled water for 30 s, then rinsed with tap water, and exposed to Eosin Y 1% alcoholic solution (5min, room temperature; Kashaf AlKawashef). Finally, the cells were washed three times with deionized water, air-dried and analysed by bright-field microscopy using the EVOS M7000 imaging system (Invitrogen). For each sample five randomly selected areas were imaged, and VSV-G protein-driven syncytium formation was quantified by measuring the total area of fusion or measuring the size of a total of 50 syncytia in each sample using ImageJ software. Syncytia were defined as cells containing at least three nuclei. To eliminate potential bias and correct for counting errors, imaging was performed independently by two blinded individuals, and the average measurements were used for each sample. Furthermore, for all biological replicates, the average syncytia size per sample was calculated from a total of 50 syncytia per sample and obtained from randomly selected areas of the well.

### Statistical Analysis

All experiments were repeated at least three times. Statistical analysis was performed using ImageJ software. The results of experimental measurements were calculated using Prism software and presented as the mean and standard error of the mean. Data was analysed statistically using paired students T-tests or one-way analysis of variance (ANOVA). A p-value of less than 0.05 was considered statistically significant. * p≤0.05, ** p≤0.01, *** p≤0.001.

## RESULTS

### UBE2J1 promotes Vesicular Stomatitis virus replication

VSV is one of the most accessible experimental models to study RNA virus infection, but has not yet been used as a model to study regulation by UBE2J1. Considering recent studies showing a role for this protein in the lifecycle and infectivity of RNA viruses such as DENV, ZIKV, SeV, H1N1 (13) and WNV (10), we investigated whether it has any impact on VSV. To this end, BHK-21 cells were transfected with an UBE2J1-expressing plasmid or an empty vector control. Twenty-four hours later the cells were infected with VSV-GFP virus (at an MOI of 1). Twenty-four hours post-infection, the culture medium was recovered and the harvested virus was titrated on a fresh monolayer of BHK-21 cells in serial dilutions so that plaque assays would allow us to assess any differences in the viral titre. The virus we used (VSV-GFP) was modified to express a GFP-tag so that it was possible to microscopically evaluate the degree of infectivity by assessing the GFP expression (Fig.1A). As shown in Fig.1 viral particles generated by UBE2J1 expressing cells showed higher levels of GFP and were associated with significantly higher levels of viral titre (PFU/ml) (Fig.1B).

**Fig. 1.**
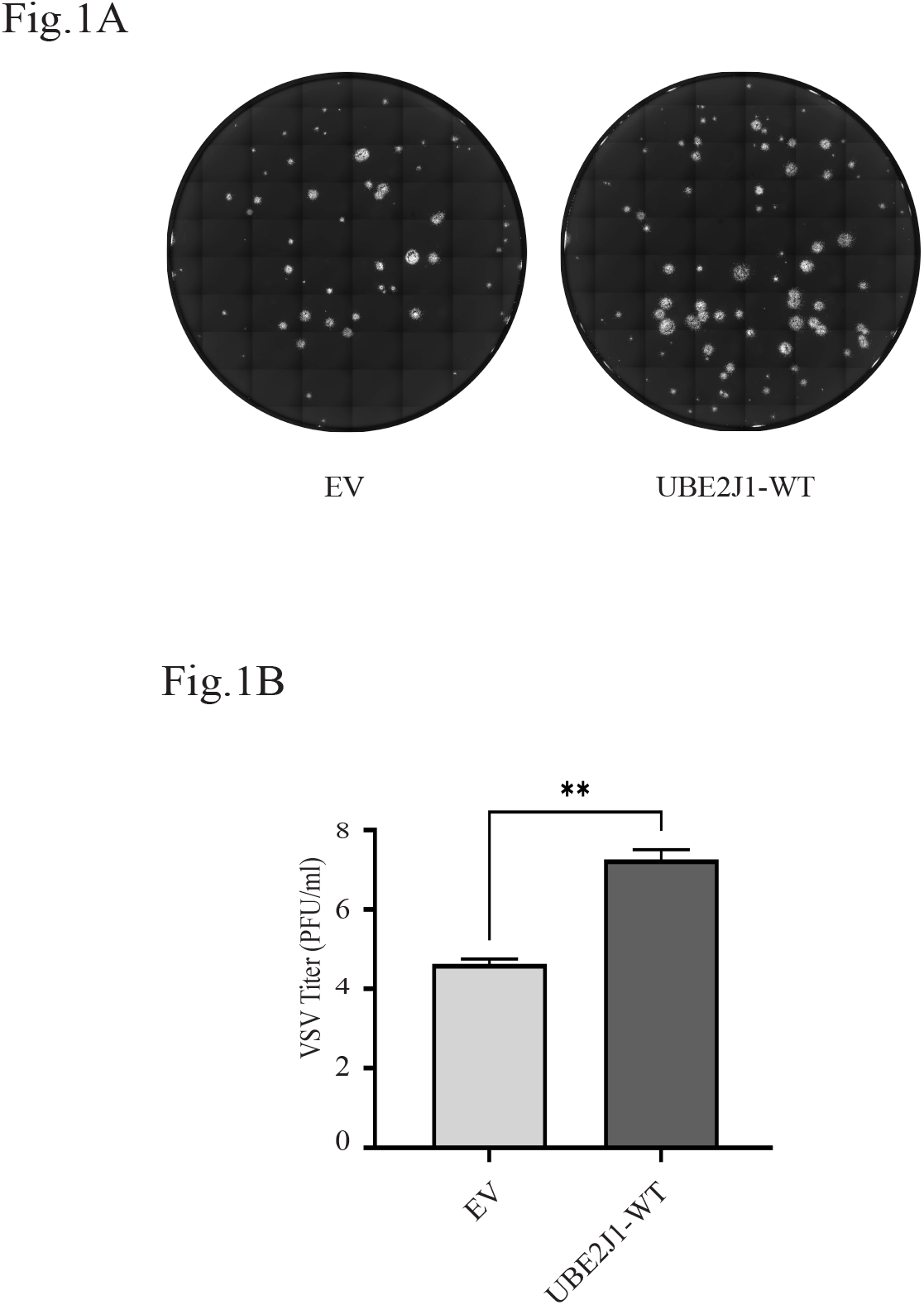
UBE2J1 promotes the replication of VSV. BHK21 cells were transiently transfected with empty vector (EV) or UBE2J1-WT-His plasmids. At 24 h post-transfection, the cells were infected with VSV-G-GFP virus (MOI of 1) for 24 h. The virus was harvested, and a plaque assay was performed for each viral sample. pc-DNA3.1 was used as an empty vector (EV) for our negative control. (A) Fluorescent microscopic examination to observe plaques and calculate PFU/mL for each plaque assay. (B) Bar graph showing PFU/mL values presented as mean ± SEM from three independent experiments. **, p ≤ 0.01.

### Co-expression of catalytically active UBE2J1 with VSV-G is associated with enhanced syncytia formation

Viral infections can result in changes in several cellular processes and parameters. While these mechanisms vary among viruses, it has specifically been shown for VSV to involve the formation of multinucleated syncytia following the membrane fusion of infected cells. Although this is not unique to VSV, its’ occurrence represents an important stage in the infectivity of the virus that facilitates replication, dissemination, and immune evasion (20). For VSV, it has been shown that syncytialization can be achieved by ectopic overexpression of the fusogenic viral VSV-G spike protein, allowing for this specific aspect of infection to be studied in the absence of other proteins from actual virus (26).

To test whether UBE2J1 contributes towards syncytialization, BHK21 cells were transfected to express VSV-G in the presence or absence of wild type UBE2J1, and after 48 h, the cells were fixed and stained. As can be seen in Fig.2A, microscopic examination showed clear differences between the transfected samples and their respective abilities to form syncytia. For cells transfected with empty vector or wild-type UBE2J1, and in the absence of VSV-G protein, there was little evidence of the development of any syncytia. However, once VSV-G was co-transfected, there was clear evidence of their formation, and UBE2J1 led to a significant increase in the fusion area (Fig.2B).

**Fig. 2.**
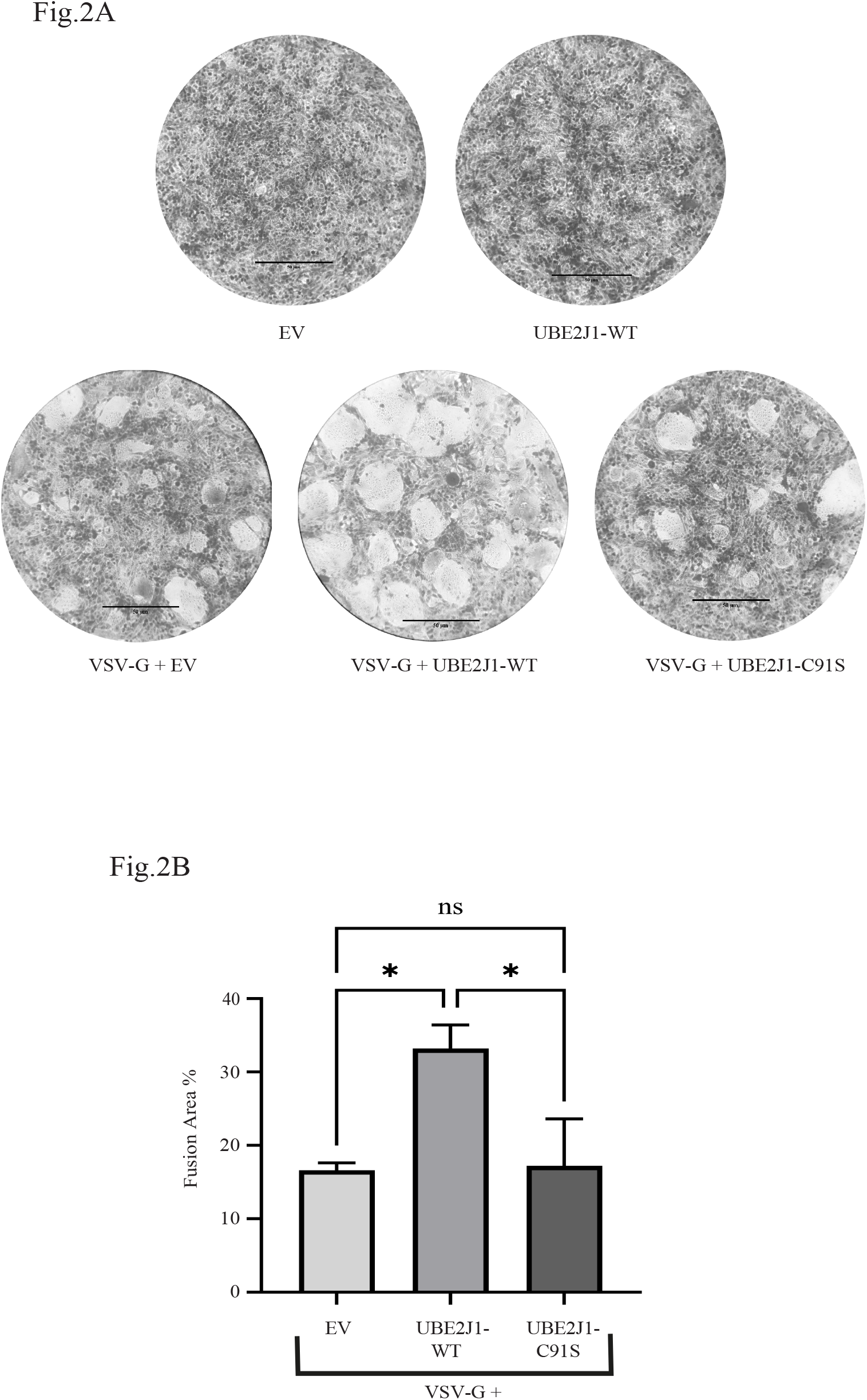
Catalytically active UBE2J1 enhances the formation of syncytia by VSV-G. BHK21 cells were transiently transfected to express wild-type (WT) or catalytically inactive (C91S) UBE2J1-His variants in the presence or absence of VSV-G, as indicated. pc-DNA3.1 was used as an empty vector (EV) for the negative controls. At 48 h post-transfection, the cells were fixed with 4% formaldehyde prior to H&E staining to reveal syncytial foci. (A) Microscopic examination was conducted to compare syncytial formation at a magnification of 20X (size bar shows 50 μm). B) Bar graph showing analysis of the fusion area % values. Data are presented as mean ± SEM from three independent experiments. **, p ≤ 0.01.

To test whether the role of UBE2J1 in syncytia formation depends on its enzymatic function, BHK21 cells transfected to express the VSV-G protein were co-transfected with enzymatically inactive UBE2J1-C91S. Following microscopic examination and quantitative analysis, it was evident that the enhancement of syncytia formation by UBE2J1 was dependent on its’ catalytic activity, and was significantly decreased by the C91S mutation (Fig. 2A and 2B).

### Phosphorylation of UBE2J1 at S184 contributes to enhanced syncytia formation

Phosphorylation of UBE2J1 at serine S184 is functionally important for cellular stress responses (7) and appears to play a role in ALV-A infection (14). To assess whether it might be of importance for syncytia formation in our transfection model, BHK21 cell monolayers expressing VSV-G were co-transfected to express either UBE2J1-WT or UBE2J1-S184A. Forty-eight hours post-transfection, H&E staining followed by microscopic examination demonstrated a clear contrast between the differentially transfected cells. As shown in Fig. 3A, and represented in the bar graph in Fig.3B, the ability of UBE2J1 to become phosphorylated at S184 is of clear importance for enhancing syncytia formation, and compared to cells expressing the wild-type UBE2J1, there was a significant decrease observed in the fusion areas for phospho-deficient UBE2J1-S184A.

**Fig. 3.**
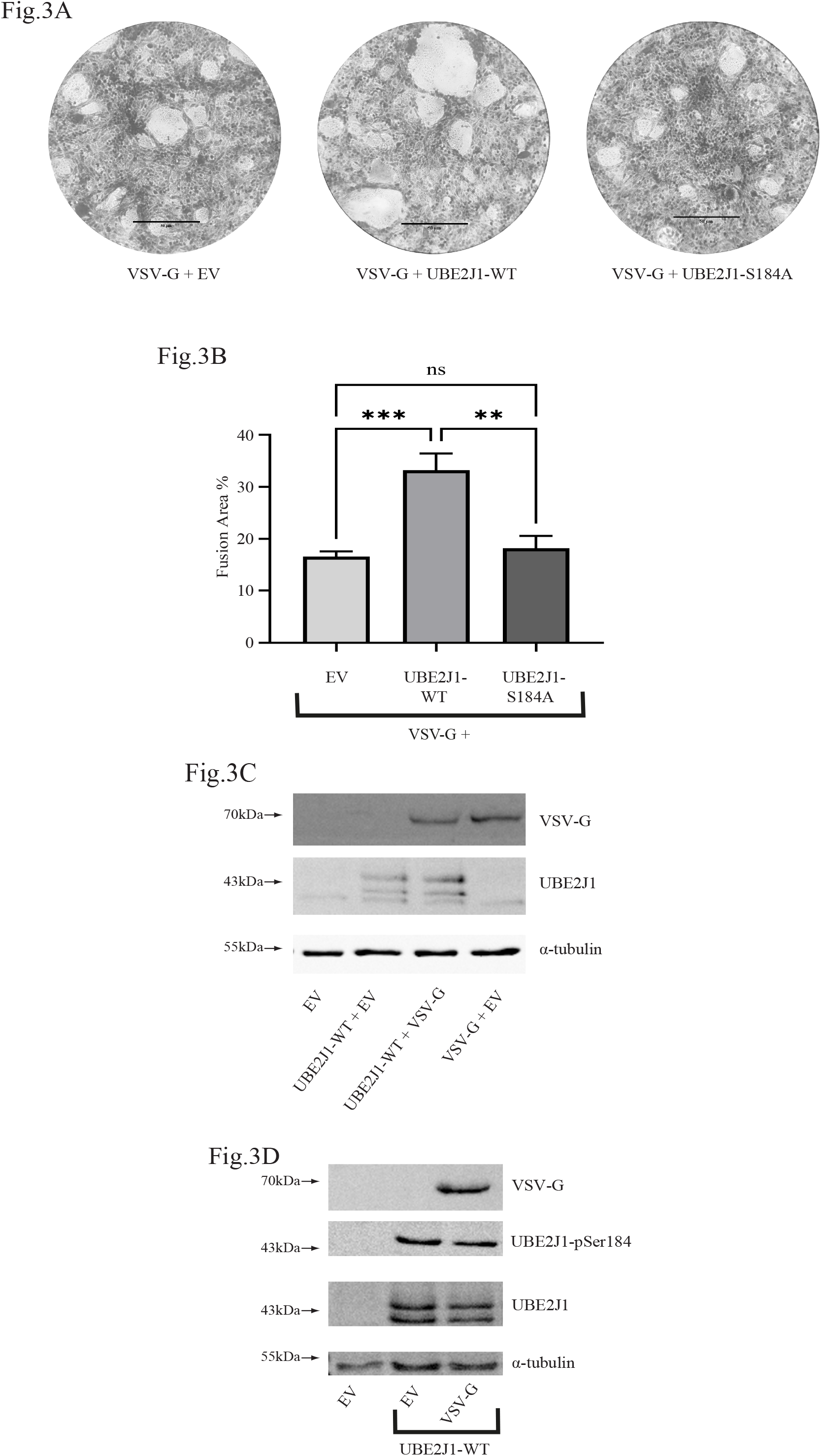
Phosphodeficient UBE2J1-S184A fails to enhance syncytia formation by VSV-G. (A and B) BHK21 cells transiently expressing VSV-G were co-transfected to express wild-type (WT) or phosphodeficient (S184A) forms of UBE2J1-His. pc-DNA3.1 was used as an empty vector (EV) control. At 48 h post transfection, the cells were fixed for H&E staining to reveal syncytial foci for microscopic examination (A) and to quantify the percentage of syncytial area per field from five fields per well (B). Data are presented as mean ± SEM of three independent experiment, ns, nonsignificant; **, p ≤ 0.01; ***, p ≤ 0.001. (C) BHK21 or (D) HEK293T cells were transfected with pcDNA (EV) or plasmids expressing UBE2J1-His or VSV-G. 48 h post transfection, harvested lysates were analysed on immunoblots using anti-His (UBE2J1), anti-phospho UBE2Jl-pSerl84, and anti-VSV-G antibodies as indicated. α-Tubulin was used as the loading control.

In light of these experimental results we wondered whether ectopic expression of the viral G-protein could actively promote the phosphorylation of UBE2J1, and explain why S184 is so important. To test this, BHK21 cells expressing UBE2J1 were co-transfected with either VSV-G or empty vector plasmids and harvested 24 h after transfection. As shown in Fig.3C, UBE2J1 was fractionated into two main bands, with the higher molecular weight form indicative of phosphorylation at S184. In the presence of VSV-G, we found no evidence to suggest a significant accumulation of the upper band and there was no detectable change in the ratio of higher to lower molecular weight forms. This was independently confirmed in lysates from transfected HEK293T cells using an anti-phospho specific antibody that preferentially recognizes phosphorylated S184 (Fig.3D).

### A cytoplasmic form of UBE2J1 increases the percent syncytial fusion area

Considering that UBE2J1 is primarily an ER localized enzyme, we wondered whether this was important for the syncytialization we observed. To test this, BHK-21 cells expressing VSV-G were co-transfected to express truncated UBE2J1-ΔTM, which lacks the transmembrane domain that anchors the carboxyl tail to the ER. Forty-eight hours post-transfection, the cells were fixed and examined microscopically. As shown in Fig.4.A, co-expression of UBE2J1-ΔTM further promoted syncytialization.

**Fig. 4.**
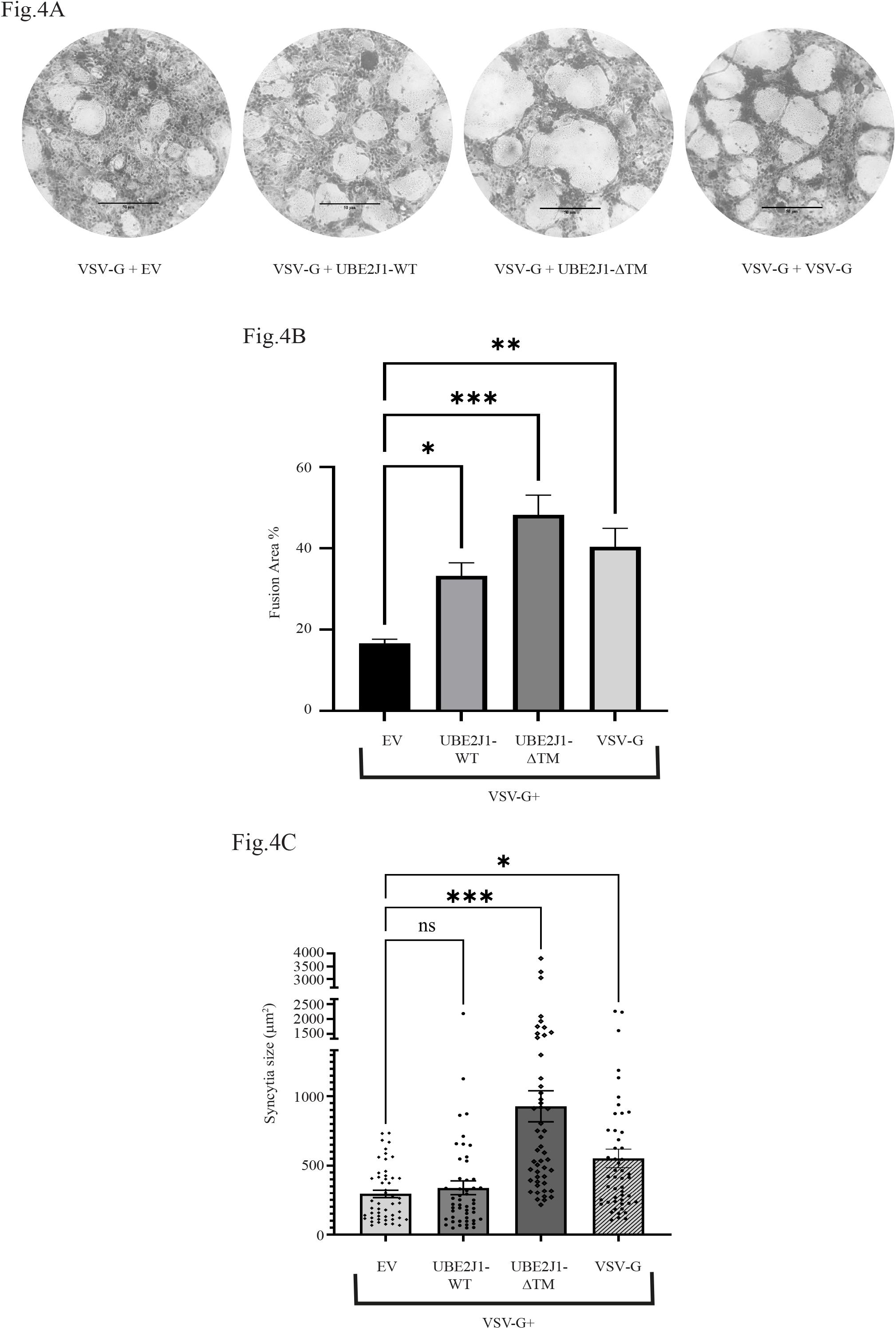
Co-expression of VSV-G with a truncated UBE2J1-ΔTM increases the size and fusion area of syncytia. BHK21 cells transiently expressing VSV-G protein were co-transfected to express wild-type UBE2J1-WT, truncated UBE2J1-Δ TM. or a double dose of VSV-G. pcDNA3.1 was used as an empty vector (EV) control. (A) At 48 h post-transfection, cells were fixed for H&E staining and light microscopy was used to compare the formation of syncytia at a magnification of 20X (size bar shows 50 μm). B) Bar graph analysis of the fusion area % values. Data arc presented as mean ± SEM from three independent experiments. (C) For each experimental sample, 50 syncytia were randomly selected from microscopic images to determine the size (μm^2^). *, p ≤0.05 **, p ≤0.01; ***,p≤0.001.

For these experiments, the fusion area for UBE2J1-ΔTM co-transfection was not only greater than that for the empty vector control, but it was also significantly larger than that for the full-length (ER membrane-embedded) form of UBE2J1 (Fig.4B). To this end, we wondered whether increasing the amount of expressed VSV-G protein would lead to an increase in syncytialization similar to that observed for UBE2J1-ΔTM. To investigate this, BHK-21 cells transfected to express the VSV-G protein were doubly co-transfected to express a second dose amount of VSV-G. Microscopic examination of stained cells after 48 h showed that doubling the amount of transfected VSV-G did indeed lead to a corresponding and significant increase in the size of the fusion area of the generated syncytia (Fig.4B). The levels achieved were comparable to those observed with the truncated UBE2J1-ΔTM.

### Expression of the carboxy-truncated UBE2J1-ΔTM protein increases the size of individual syncytia

To this point, our experiments demonstrated that UBE2J1 promotes the syncytial fusion area associated with VSV-G protein expression, with truncated UBE2J1-ΔTM leading to the greatest increase. Considering that the fusion area is based on both the number of individual syncytia and their sizes (26), we wished to differentiate between these two parameters and provide greater insight into the role of UBE2J1. BHK-21 cells were transfected to express the VSV-G protein and co-transfected with either wild-type UBE2J1, truncated UBE2J1-ΔTM or a second dose of VSV-G. Forty-eight hours post-transfection and after H&E staining, random microscopic images were captured from each transfection group to measure the sizes of 50 syncytia per sample. The sizes of individual syncytia were plotted against the different co-transfected plasmids, and as shown in Fig.4C, co-transfection of truncated UBE2J1-ΔTM with VSV-G resulted in syncytia with significantly larger size.

### The influence of UBE2J1 variants on syncytialization by the VSV-G protein is exactly mirrored in the replicative abilities of VSV

It is generally accepted that the formation of syncytia is linked to increased infectivity. Having identified mutant (C91A and S184A) and truncated (ΔTM) forms of UBE2J1 that could impact syncytia formation, we explored whether they could also influence infection with the actual virus. To investigate this, BHK-21 cells were transfected to express wild-type, truncated and mutated forms of the protein. Twenty-four hours later, the cells were infected with VSV-GFP virus (MOI of 1). Twenty-four hours post-infection, the supernatants from infected cells were harvested and viral titres were determined for each virus sample using plaque assays.

As shown by the presence of GFP-tagged plaques in Fig.5, both wild-type and ΔTM proteins that enhanced syncytialization were associated with higher infectivity, and showed more plaques than the control (Fig.5A). Thus, there was a clear association between syncytialization and viral titres, which was confirmed by calculating the PFU/mL for each sample (Fig.5B). For the full-length protein, this was dependent on enzyme activity, and the catalytically inactive C91A mutant failed to show any such enhancement. Similarly, without the capacity for phosphorylation at serine residue S184, there was no evidence to indicate that the UBE2J1-S184A mutant could enhance infectivity.

**Fig. 5.**
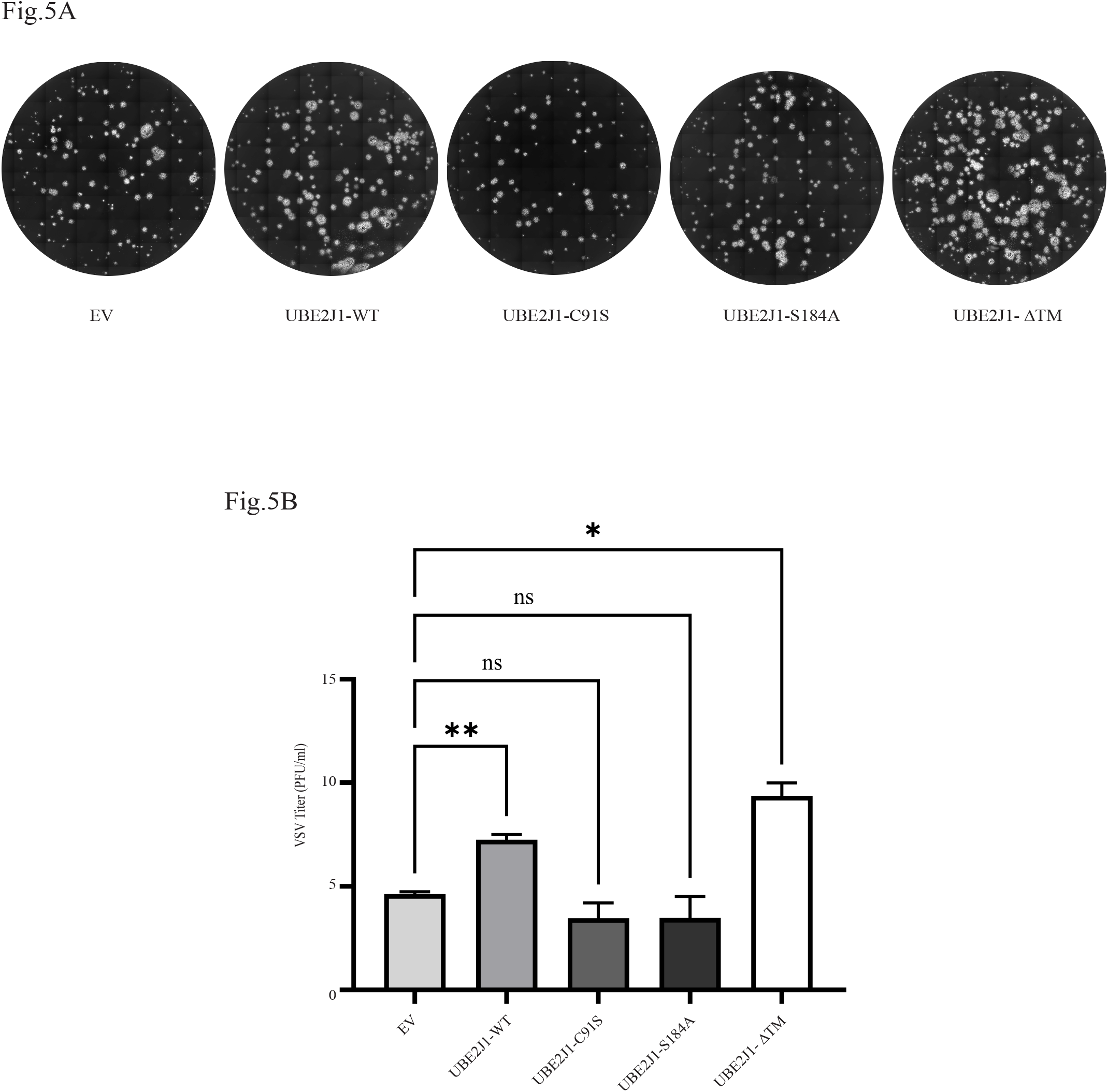
UBE2J1 variants impact on VSV infectivity. BHK21 cells were transiently transfected to express wild-type UBE2J1-WT-His, catalytically inactive UBE2J1-C91S-His, phosphodeficient UBE2J1-S184A-His, or carboxy-truncated UBE2J1-ΔTM-His. pcDNA3.1 was used as negative control (EV). At 24 h post-transfection, cells were infected with VSV-G-GFP virus (MOI of 1) for 24 h. The virus was harvested, and plaque assay was performed for each sample. (A) Fluorescent microscopic examination of infected cells was used to compare the plaques and (B) to calculate the PFU/mL for each plaque assay. Data are presented as mean ± SEM from three independent experiments. ns, nonsignificant; *, p ≤ 0.05; **, p ≤ 0.01.

## DISCUSSION

Although UBE2J1 is best recognised for degrading misfolded proteins as a result of environmental stress or genetic mutations (27), it also contributes to the proteasomal degradation of viral proteins in preparation for MHC class I presentation to cytotoxic T cells (2, 3, 8). In this respect it contributes to the effective clearance of viruses. However recent evidence indicates that UBE2J1function can also be subverted to support and promote viral infection. This includes the disruption of signalling pathways that influence interferon production (13-15), with consequences for several aspects of viral infection and the immune response. The present study aimed to assess the involvement of UBE2J1 in VSV infection, which is a non-segmented, negative-sense RNA virus.

Our studies from the outset indicated a role for UBE2J1 in propagating the infection, with results pointing towards a specific enhancement of syncytialization associated with ectopic expression of the VSV-G protein. Considering that UBE2J1 is part of the ERAD translocon, it remains a possibility that our observations reflect a structural role in a larger multiprotein complex (2, 3) rather than catalytic function. However, mutation of the cysteine residue in the active site abolished this enhancement, confirming the importance of the enzyme’s catalytic activity. UBE2J1 is known to suppress the antiviral response by promoting the ubiquitination and degradation of both IRF3 (13) and IRF7 (15), and by inhibiting the STAT3/IRF1 pathway (14). Disruption of the IRF signalling network can reduce anti-viral interferon production, which would result in increased levels of VSV-G (28). However, given that interferon signalling involves the regulation of Interferon Induced TransMembrane (IFITM) proteins, which can act to suppress viral entry, there may be more specific mechanistic links to syncytia formation (18). Of these, IFITM1 is known to play a particularly important role, and certainly in the case of VSV, IFITM3 has been shown to suppress VSV-G mediated viral entry (29). Therefore, one possible explanation for our results is that suppression of the IRF3, IRF7, and STAT3/IRF1 signalling network leads to decreased cellular IFITM levels, with an associated increase in syncytialization.

Our studies not only show that UBE2J1 enhances viral infection and syncytia formation, but also highlight a role for phosphorylation of the protein at S184. Phosphorylation of UBE2J1 is important for cellular responses to misfolded proteins and is known to influence interactions with E3 ubiquitin ligase enzymes (7). While there is some evidence that phosphorylation influences catalytic activity (4), the E3 ligases that cooperate with UBE2J1 to regulate the interferon response have not yet been fully identified and a possible role for phosphorylation in regulating either interferon signalling or syncytialization remains poorly understood. Interestingly, Wang *et al*. have shown a role for UBE2J1 in ALV-A viral replication, with an S184A mutant exhibiting less suppression of STAT3/IRF1 signalling compared to the wild-type protein (14). However, that study did not specifically address syncytia formation by ALV-A, and whether UBE2J1 might be influence this specific aspect of infection.

The functional role of UBE2J1 is linked to its’ localisation at the ER membrane and its’ contribution to the translocon (4). Given that syncytia formation depends on lipid membrane restructuring and fusion (20, 30, 31) we hypothesized that the enhancement that we observed would require membrane anchoring of UBE2J1. Instead, we found that the soluble cytoplasmic form of the protein had the greatest effect, resulting in larger syncytia formation. In the context of previous studies, a dose-response relationship exists between VSV-G and syncytia formation, with the inference that increasing infection titre leads to higher VSV-G protein levels, which in turn promotes greater syncytialization and viral propagation (32) In our study, we demonstrated that doubling the amount of transfected VSV-G plasmid into the cells increased both the syncytia fusion area and the average size of individual syncytia. However, the contribution of truncated UBE2J1-ΔTM to syncytia formation exceeded even the effect of doubling the transfected VSV-G plasmid. This suggests that cytoplasmic localisation of UBE2J1 could be of particular importance in enhancing VSV infection. This is notable in the context of previous reports showing that EV71 proteases can cleave UBE2J1 at Q273 (33) (which is proximal to residue 287 at which our ΔTM is truncated). The Wang et al. study emphasized that this could represent a novel mechanism for the virus to disrupt ERAD (33). However, it did not account for the possibility that cleavage could result in the cytoplasmic accumulation of a catalytically active form of UBE2J1 (33). Although our studies did not specifically address whether VSV promotes the proteolysis of UBE2J1, and unlike EV71, the VSV genome does not code for any proteases, the expression of the truncated form of the protein resulted in a clear increase in syncytia formation. For viruses that encode proteases that can cleave UBE2J1, this could represent further subversion of protein function to promote infection.

Although the exact molecular links between syncytia formation and infectivity are still being elucidated (34), and the molecular factors that regulate the sizes of individual syncytia are not well understood, we completed our study by examining the impact of the different UBE2J1 protein isoforms on viral titres. These experiments showed a decrease for the C91S and S184A mutants and a clear increase in viral replication and infectivity for the truncated UBE2J1-ΔTM, supporting a mechanistic link with larger syncytia sizes. In conclusion, our study demonstrated the involvement for UBE2J1 in VSV replication with enhanced syncytialization associated with more severe infection.

## ACKNOWLEDGEMENTS

The authors wish to thank the Ministry of Education, Saudi Arabia for supporting N.D Algoufi, and T.M.Wac for useful suggestions.

## ABBREVIATIONS

Ubc: ubiquitin conjugating
UPR: Unfolded Protein Response
ERAD: endoplasmic reticulum associated degradation
VSV: Vesicular Stomatitis Virus

