## Supplementary Figure for "Phosphorylation of UBE2J1 at serine residue S184 contributes towards infection and cellular syncytialization by Vesicular Stomatitis Virus"

### Supplementary Materials

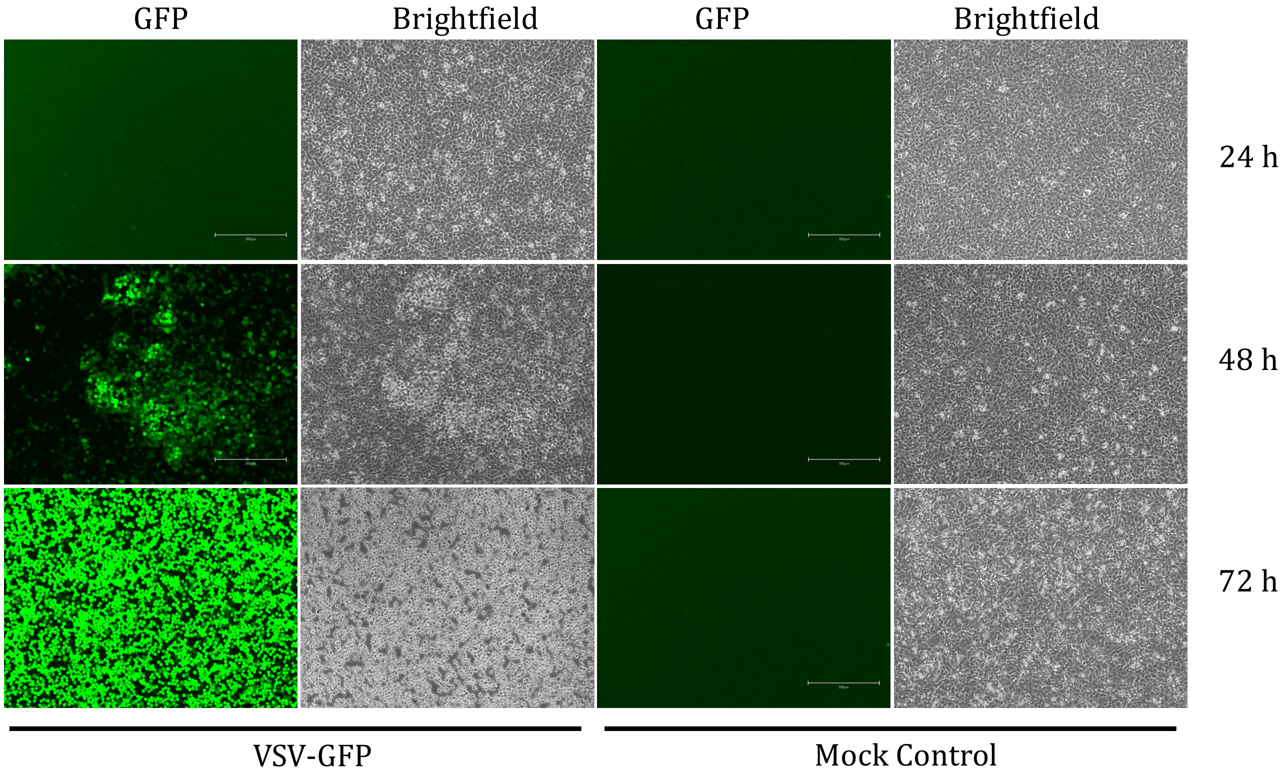


**Supplementary Figure. Rescue of rVSV-GFP virus.**
BHK-21 cells seeded in 6-well-plates were infected with filtered supernatant and monitored for 3 days for the development of VSV-specific CPEs and GFP expression. After 72 hours, cells showed very strong VSV-specific CPE represented by the rounding and detachment of cells and very clear GFP indicating the successful virus recovery. Rescued virus was then propagated on BHK-21cells to reach very high titre.
